# Complex and critical roles for the AtrR transcription factor in control of *cyp51A* expression in *Aspergillus fumigatus*

**DOI:** 10.1101/2022.01.28.478276

**Authors:** Sanjoy Paul, Paul E. Verweij, Willem J.G. Melchers, W. Scott Moye-Rowley

## Abstract

*Aspergillus fumigatus* is the major filamentous fungal pathogen in humans. The gold standard treatment of *A. fumigatus* is based on azole drug use but the appearance of azole-resistant isolates is increasing at an alarming rate. The *cyp51A* gene encodes the enzymatic target of azole drugs and azole-resistant alleles of *cyp51A* often have an unusual genetic structure containing a duplication of a 34 or 46 bp region in the promoter causing enhanced gene transcription. These tandem repeats are called TR34/TR46 and produce duplicated binding sites for the SrbA and AtrR transcription factors. Using site-directed mutagenesis, we demonstrate that both the SrbA (sterol response element: SRE) and AtrR binding sites (AtrR response element: ATRE) are required for normal *cyp51A* gene expression. Loss of either the SRE or ATRE from the distal 34 bp repeat of the TR34 promoter (further 5’ from the transcription start site) caused loss of expression of *cyp51A* and decreased voriconazole resistance. Surprisingly, loss of these same binding sites from the proximal 34 or 46 bp repeat led to increased *cyp51A* expression and voriconazole resistance. These data indicate that these duplicated regions in the *cyp51A* promoter function differently. Our findings suggest that the proximal 34 or 46 bp repeat in *cyp51A* recruits a corepressor that requires multiple factors to act while the distal repeat is free of this repression and provides the elevated *cyp51A* expression caused by these promoter duplications.

**Importance:** *Aspergillus fumigatus* is the most common human filamentous fungal pathogen. Azole drugs are the current therapy of choice for *A. fumigatus* but the prevalence of azole resistance is increasing. The main genetic alteration causing azole resistance is a change in the *cyp51A* gene that encodes the target of these drugs. Azole-resistant *cyp51A* alleles routinely contain duplications in their promoter regions that cause increased gene transcription. Here, we demonstrate that clinical isolates containing a 34 or 46 bp duplication in the *cyp51A* promoter required the presence of the transcription factor-encoding *atrR* gene to exhibit elevated azole resistance. Elimination of transcription factor binding sites in the *cyp51A* gene have differential actions on expression of the resulting mutant allele. These data dissect the molecular inputs to *cyp51A* transcription and reveal a complicated function of the promoter of this gene that is critical in azole resistance.

## Results and discussion

*Aspergillus fumigatus* is the most common cause of mold infections in humans (1). Azole drugs are currently the first-line therapy for aspergillosis. However, azole-resistant *A. fumigatus* clinical isolates are being found with increasing frequency and are associated with a significantly worse clinical outcome (2). Although multiple mechanisms contribute to azole resistance in *A. fumigatus*, the most commonly reported genetic change associated with this phenotype are alterations in the gene encoding *cyp51A*, the target enzyme of azole drugs (3). The most prevalent azole resistance allele is a compound mutation in *cyp51A* consisting of a 34 bp duplication in the promoter element (TR34) and a single amino acid replacement in the coding sequence (L98H) (4). Both of these mutations are required for the observed high level azole resistance conferred by this compound allele (5).

Although it is well-established that the TR34 *cyp51A* promoter drives increased expression of *cyp51A* mRNA compared to the wild-type version (5), we lack a detailed understanding of how this increased expression is achieved. Previous studies from our lab and others have demonstrated that several different transcription factors control transcription of *cyp51A* via the 34 bp region. First, the sterol-responsive SrbA regulator binds to an element in this 34 bp region called the sterol response element (SRE) and stimulates expression when sterols are limiting (6, 7). Second, the AtrR transcription factor binds to a second site within the 34 bp region, referred to as the AtrR response element (ATRE), to activate transcription (8, 9). Finally, two different negative transcriptional regulators repress *cyp51A* expression. The CCAAT-binding complex (CBC) or the iron-responsive transcription factor HapX (10) both reduce *cyp51A* expression: CBC binds within the 34 bp region while HapX binds just 3’ to this segment (11, 12). The locations of these sites and their positions relative to the 34 bp region are shown in Supplementary Figure 1A. Note that both the TR34 and TR46 promoters contains two SREs and ATREs owing to the 34 bp duplication. The CBC binding site is also duplicated but the HapX response element (HXRE) is not. To distinguish between these two copies of each site, we refer to them as either the proximal SRE/ATRE (proximal; closest to transcription start, pSRE/pATRE) or distal SRE/ATRE (distal; furthest from transcription start, dSRE/dATRE).

To evaluate how the ATRE, SRE and HXRE contribute to expression of both wild-type and TR34 versions of *cyp51A*, site-directed mutations were constructed in these elements (Supplementary Figure 1A) and returned to the natural *cyp51A* genomic location (Figure 1A). These strains were tested for their ability to grow in the presence of voriconazole (Figure 1B) and the level of *cyp51A* expression was evaluated by reverse transcription-quantitative PCR (RT-qPCR) (Figure 1C) or using an anti-Cyp51A antibody (Supplementary Figure 1B) analyses.

**Figure 1.**
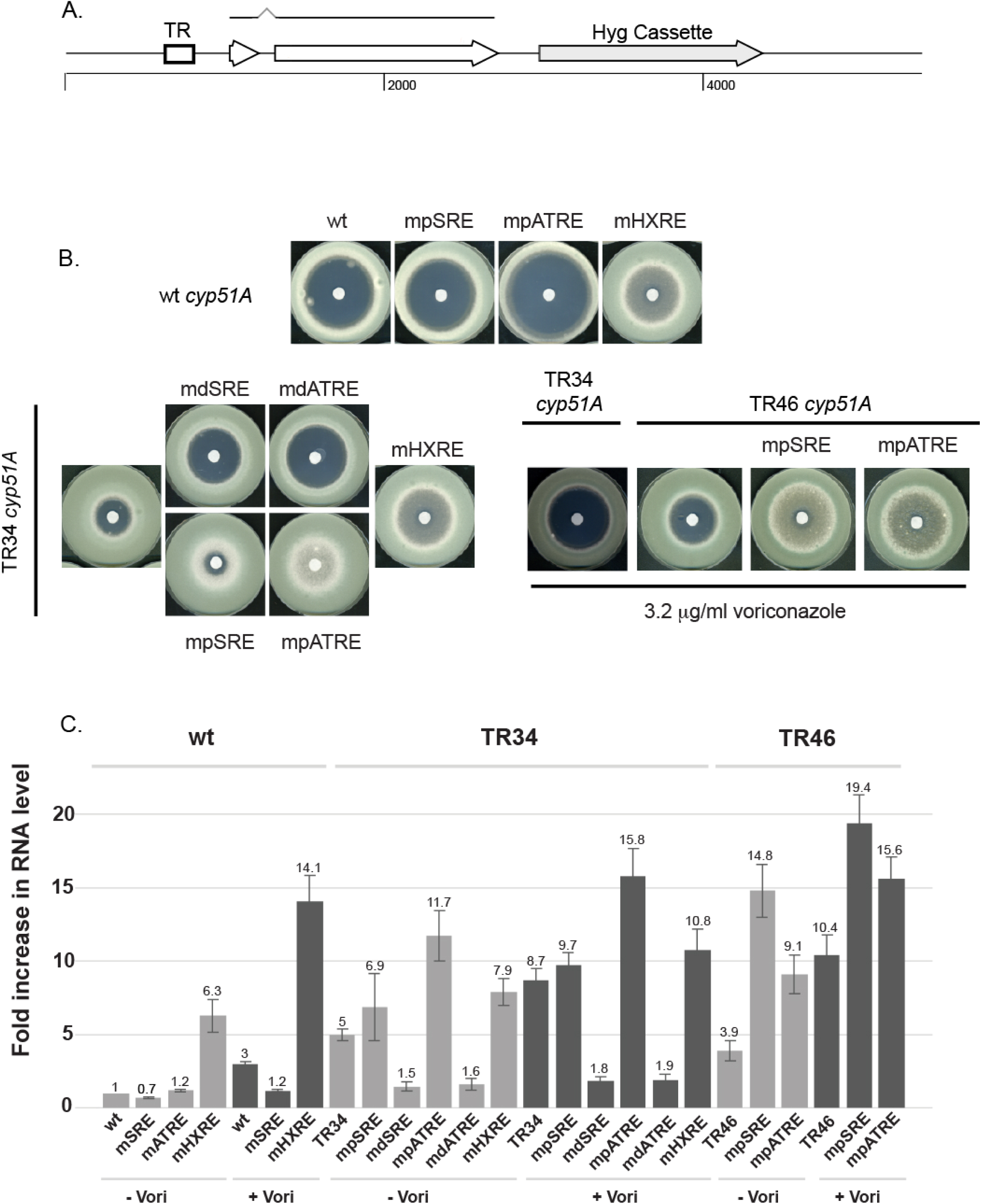
Analysis of *cyp51A* promoter function. A. Schematic diagram of integration of wild-type and mutant *cyp51A* promoter mutants. All mutants analyzed here were reintroduced back at the native *cyp51A* chromosomal location. The relative location of the TR repeat regions is indicated as a box with the two exons of *cyp51A* also noted. The hygromycin selection marker (Hyg cassette) is located downstream of the native 3’ end of the *cyp51A* mRNA. B. Disk diffusion assay of mutant forms of the *cyp51A* promoter. A filter disk containing 0.01 mg of voriconazole was placed in the center of 10^6^ spores of each indicated strain and allowed to grow at 37° C for 72 hrs. C. Strains containing the listed versions of either the wild-type (left hand side) or TR34 (right hand side) *cyp51A* gene were grown to mid-log phase with (+) or without (-) voriconazole treatment. Transcriptional behavior of each mutant promoter was assessed by qRT-PCR relative to the *tef1* gene. Data are presented for the average of two independent experiments. Numbers above each bar represent the average fold increase for each strain in the presence or absence of voriconazole.

Loss of either the pSRE (mpSRE) or the pATRE (mpATRE) from wild-type *cyp51A* caused a slight (mpSRE) or a large increase (mpATRE) in voriconazole susceptibility (Figure 1B). Removal of the HXRE led to a large decrease in voriconazole susceptibility. These resistance data were fully consistent with the observed expression changes seen by either RT-qPCR measurements (Figure 1C) or western blotting (Supplementary Figure 1B). Loss of the ATRE from the wild-type *cyp51A* promoter caused such profound hypersensitivity to voriconazole that we were unable to recover sufficient fungus to assay expression. Together, these data are consistent with both the SRE and ATRE acting as positive regulatory elements and the HXRE acting as a negative element to control *cyp51A* expression and function.

Insertion of the TR34 promoter into the *cyp51A* locus led to a decrease in voriconazole susceptibility as seen before (4, 13). Strikingly, loss of either the pSRE or the pATRE from TR34 *cyp51A* led to a large decrease in voriconazole susceptibility (Figure 1B). This decrease in voriconazole susceptibility was accompanied by a large increase in the level of Cyp51A expression (Figure 1C, Supplementary Figure 1B). Behavior of each of these proximal element mutations was similar to that caused by loss of the HXRE from TR34 *cyp51A*. Although these proximal binding sites clearly work as primarily as positive elements in the wild-type promoter context, they appear to be involved in repression in the TR34 promoter as their loss leads to a large increase in *cyp51A* expression. Conversely, loss of either of the distal elements (dSRE or dATRE) caused an increase in voriconazole susceptibility, along with a decrease in expression and loss of voriconazole inducibility of *cyp51A* mRNA (Figure 1C) consistent with these binding sites acting as positive sites determining TR34 promoter function.

We also produced proximal ATRE and SRE mutant forms of the TR46 cyp51A gene to determine if the unexpected behavior of these elements would extend to this different promoter context. TR46 corresponds to duplication of 46 bases with an identical 5’ endpoint to TR34 and additional 12 bp at the 3’ end (14). As seen for their counterparts in the TR34 promoter, loss of either the proximal SRE or ATRE caused a decrease in voriconazole susceptibility and an increase in expression compared to the starting TR46 promoter-containing strain.

These data indicate that the increased Cyp51A expression and reduced voriconazole susceptibility caused by the TR34 or TR46 promoters cannot be explained simply by the increased dosage of the duplicated regions present. The proximal and distal regions have distinct behaviors in the TR34 promoter context and likely in the TR46 promoter as well. The distal 34 bp region behaves strictly as a positive regulator of *cyp51A* transcription while the proximal element exhibits a negative effect when present in the TR34 promoter. This is quite surprising since loss of the pATRE from the wild-type *cyp51A* promoter yields a strain that cannot grow in the presence of voriconazole. These same behaviors are seen for the pSRE, although this strain grew, albeit slowly, in the presence of voriconazole.

Given the important role of AtrR in control of *cyp51A* promoter function, we compared the requirement for this factor in voriconazole resistance and Cyp51A expression in wild-type and isogenic TR34 *cyp51A* laboratory strains. We also examined the effect of loss of AtrR in two different clinical strains containing either a TR34 promoter-driven *cyp51A* gene or a TR46 *cyp51A* locus. Each of the clinical isolates tested is associated with different mutant forms of Cyp51A. The *atrR* gene was disrupted in all these 4 strains using CRISPR/cas9 and isogenic *atrR* and *atrRΔ* derivatives tested for voriconazole susceptibility (Figure 2A) and expression of Cyp51A by western blotting (Figure 2B).

**Figure 2.**
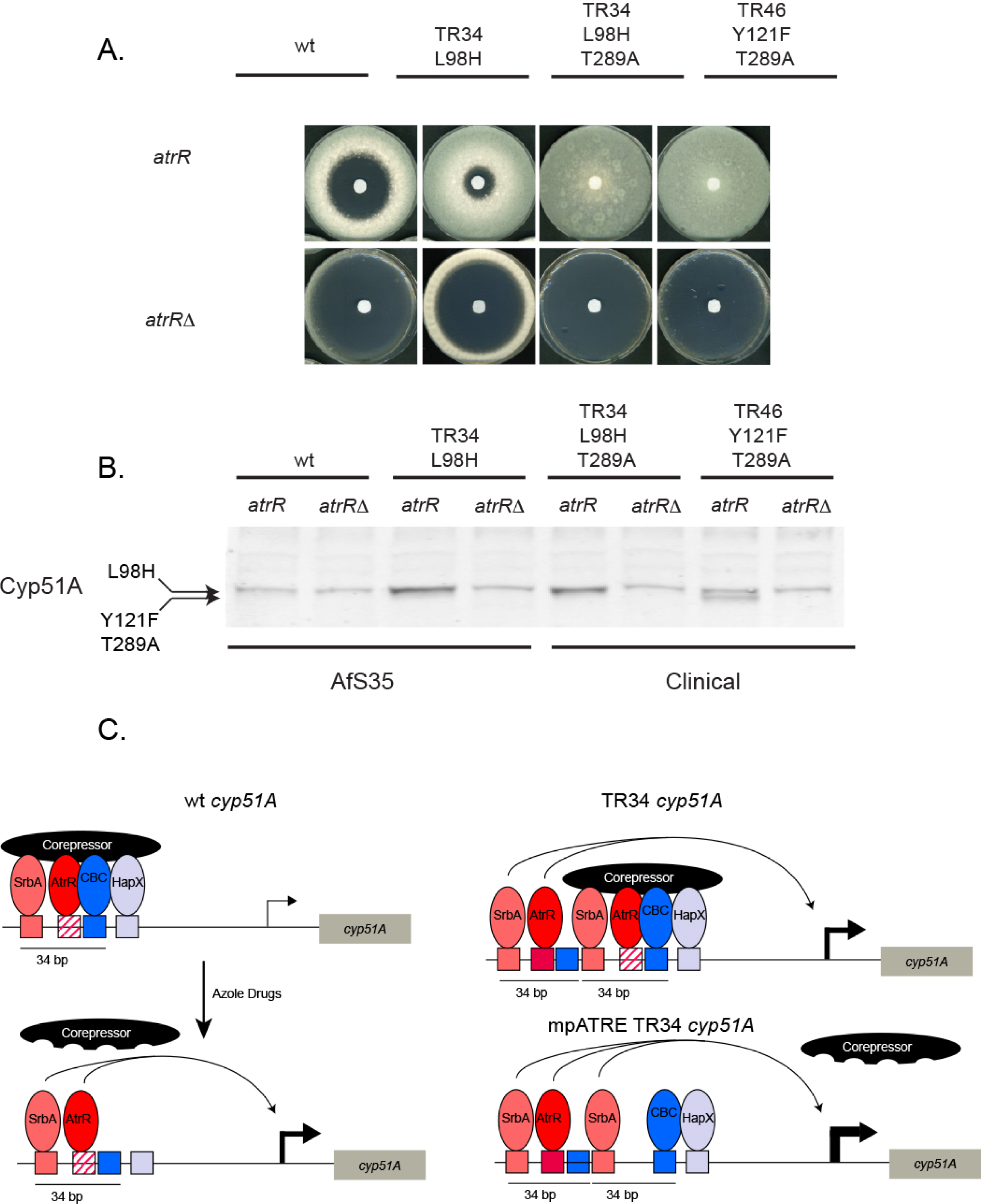
AtrR is essential for voriconazole resistance in laboratory and clinical strains. A. Isogenic *atrR* and *atrRΔ* derivatives of the indicated strains were tested for voriconazole resistance by disk diffusion assay. B. Western blot analysis of the strains listed above was performed using the anti-Cyp51A antiserum. Note that the L98H-containing enzymes electrophorese very close to a nonspecific background signal as we showed earlier (13). The Y121F T289A Cyp51A is clearly resolved below this background polypeptide. C. Diagram for potential roles of trans- and cis-acting factors at wild-type and TR34 *cyp51A* promoters. A hypothetical corepressor is pictured that makes multivalent contacts with the key regulators of *cyp51A* transcription. Note the proximal ATRE is indicated as the red crosshatched box. Other binding sites are color-coded with their respective regulators. Azole drugs trigger corepressor dissociation and gene activation. In the case of the TR34 promoter (right hand panels), the distal SRE and ATRE in the upstream 34 bp repeat can bypass corepressor function and activate transcription. The 34 (and 46) bp tandem repeats do not include the HXRE but maintain a CBC binding site. Interaction of CBC with the adjacent HXRE is required for strong binding of these factors (17). Exposure of the TR34 *cyp51A* gene to azole drugs or loss of the pSRE or pATRE (shown here) trigger strong induction of expression. Induction of expression in the mpATRE TR34 promoter is maximal even in the absence of azole induction. Only TR34 is shown but we believe the same mechanisms operate for the TR46 promoter.

The presence of AtrR was essential for the normal high level voriconazole resistance seen in both clinical isolates, irrespective of the TR34 or TR46 nature of the *cyp51A* promoter. The overexpression of Cyp51A was also eliminated from these strains when the *atrR* was deleted.

Together, these data illustrate the unexpected complexity of the TR34 promoter region in *cyp51A* expression. We argue that a simple increase in dosage of a positively acting region of 34 bp cannot explain the unique behavior of the TR34 promoter. The distal 34 bp repeat behaves positively but the proximal 34 bp repeat has a strong negative effect on TR34 promoter activity. We hypothesize the presence of a multivalent corepressor (Figure 2C) that must be engaged by SrbA and AtrR, along with CBC and HapX to normally repress *cyp51A* transcription. A single transcription factor acting as both a repressor or activator has been extensively documented for mammalian nuclear receptors (15). Loss of the binding sites for SrbA or AtrR strongly activate *cyp51A* expression in the absence of drug induction but only in the context of a duplication of the cyp51A promoter. Importantly, neither the TR34 or TR46 duplication includes both the CBC and HapX binding sites, suggesting that these must be lost in order to provide the proper context for the upstream repeat to induce *cyp51A* expression. In the wild-type *cyp51A* promoter, mutations in either the SRE or the ATRE cannot hyperactivate since these elements are also required for normal expression. AtrR is required for voriconazole resistance and Cyp51A overproduction from TR34 and TR46 promoter-driven *cyp51A* genes and, as seen earlier with SrbA (16), AtrR is a crucial determinant for azole resistance in clinical isolates of *A. fumigatus*.

## Supplementary Figures and Material and Methods

**Supplementary Figure 1.**
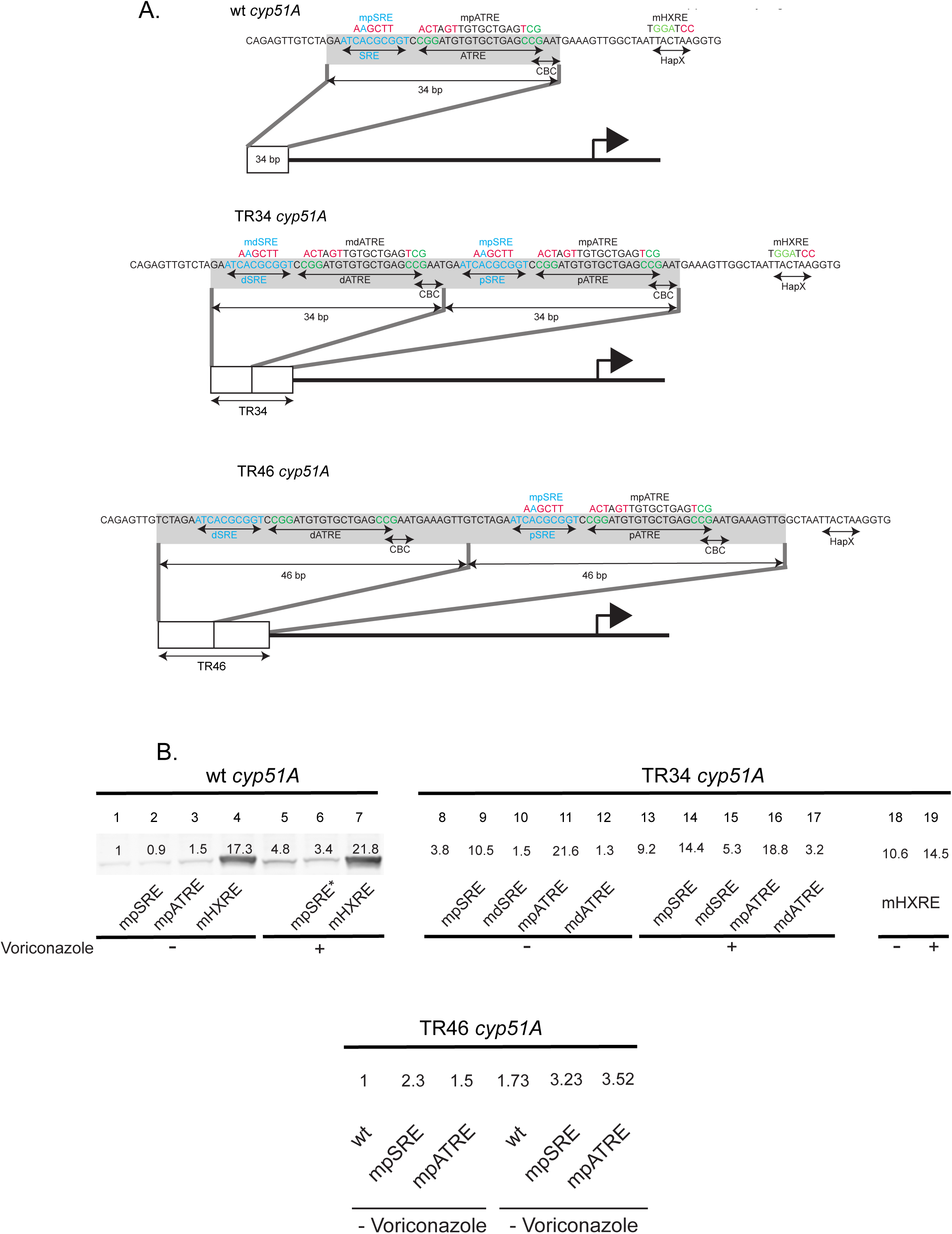
Detailed map of *cyp51A* promoter mutations and analysis of Cyp51A protein levels in response to these alterations. A. The DNA sequence of the cyp51A promoter region of interest in this study is shown. The wild-type promoter is shown on the top and the TR34 equivalent is shown on the bottom. Location of the core binding elements for each transcription are indicated below the DNA sequences. Mutant bases are shown in red lettering in each site. Extent of the 34 bp repeat is shown by the gray highlighting. B. Whole cell protein extracts were prepared and analyzed by western blotting using the anti-Cyp51A antiserum (13). Strains lacking the pATRE in the wild-type *cyp51A* promoter context were unable to be grown in the presence of voriconazole and are absent from that analysis. Lanes are numbered at the top of each panel and the numbers near each Cyp51A polypeptide correspond to the quantitation for this experiment.

## Materials and Methods

### Strains & growth conditions

The lab strains used in this study were derived from the AfS35 (FGSC #A1159). Strains used here are listed in Table 1. *A. fumigatus* strains were typically grown at 37°C in rich medium (Sabouraud dextrose: 0.5% tryptone, 0.5% peptone, 2% dextrose [pH 5.6]). Selection of transformants and the drug disc diffusion assay was performed in minimal medium (MM: 1% glucose, nitrate salts, trace elements, 2% agar [pH 6.5]); trace elements, vitamins, and nitrate salts, supplemented with 1% sorbitol and either 200 mg/liter hygromycin Gold (InvivoGen) or 0.1 mg/litre pyrithiamine. For solid medium, 1.5% agar was added.

**TABLE 1.** *A. fumigatus* strains used in this study

| Strain | Parent | Genotype | Source or reference |
| --- | --- | --- | --- |
| V232-12 |  | TR34 L98H T289A *cyp51A* | W. Melchers |
| SPF169 | V232-12 | *atrRΔ::ptrA* | This study |
| V181-51 |  | TR46 Y121F T289A *cyp51A* | W. Melchers |
| SPF169 | V181-51 | *atrRΔ::ptrA* | This study |
| AfS35 | D141 | *akuA::loxP* | FGSC |
| SPF92 | AfS35 | wt *cyp51A* hph | (13) |
| SPF200 | AfS35 | mSRE *cyp51A* hph | This study |
| SPF202 | AfS35 | mATRE *cyp51A* hph | This study |
| SPF204 | AfS35 | mHXRE *cyp51A* hph | This study |
| SPF94 | AfS35 | TR34 *cyp51A* hph | (13) |
| SPF206 | AfS35 | mdSRE TR34 *cyp51A* hph | This study |
| SPF208 | AfS35 | mdATRE TR34 *cyp51A* hph | This study |
| SPF210 | AfS35 | mHXRE TR34 *cyp51A* hph | This study |
| SPF212 | AfS35 | mpSRE TR34 *cyp51A* hph | This study |
| SPF214 | AfS35 | mpATRE TR34 *cyp51A* hph | This study |

### Transformation and generation of *A. fumigatus cyp51A* mutants

The plasmid backbone for generating *A. fumigatus cyp51A* promoter mutants was provided by Eveline Snelders and colleagues. This plasmid has a wild-type *cyp51A* promoter, gene and transcription terminator followed by a hygromycin resistance cassette (hph) and 1.3 kb downstream of *cyp51A* for homologous targeted integration (5). The specific *cyp51A* promoter mutations were synthesized as DNA fragments and obtained from GenScript USA Inc. The DNA fragments were then subcloned into the vector backbone by digesting the vector and synthesized DNA fragments with PmlI and BglII. SRE, ATRE and HXRE mutations were marked with HindIII, SpeI and BamH1 restriction sites, respectively. The recombinant plasmids were verified by PCR amplifying a 1kb region around the mutation sites, and restriction digests using the appropriate restriction enzymes. The recombinant plasmids carrying the different *cyp51A* promoters were digested with PmlI and PstI to release the *cyp51A* promoter-gene-hph cassette for transformation. The list of plasmids used for this study is listed in Table 2.

**TABLE 2.** Plasmids used in this study

| Plasmid | Parent | Genotype | Source or reference |
| --- | --- | --- | --- |
| A1 | pUC57 | wt *cyp51A* hph | (5) |
| pSP119 | A1 | mSRE *cyp51A* hph | This study |
| pSP120 | A1 | mATRE *cyp51A* hph | This study |
| pSP121 | A1 | mHXRE *cyp51A* hph | This study |
| L5H | pUC57 | TR34 *cyp51A* hph | (5) |
| pSP122 | A1 | mdSRE TR34 *cyp51A* hph | This study |
| pSP123 | A1 | mdATRE TR34 *cyp51A* hph | This study |
| pSP124 | A1 | mHXRE TR34 *cyp51A* hph | This study |
| SPF125 | A1 | mpSRE TR34 *cyp51A* hph | This study |
| SPF126 | A1 | mpATRE TR34 *cyp51A* hph | This study |

Transformation was performed using in vitro-assembled cas9-guide RNA-ribonucleoproteins coupled with 50 bp microhomology repair templates (18). Transformants with targeted integration were confirmed by diagnostic PCR of the novel downstream junction as well as by PCR amplification and subsequent sequencing of the *cyp51A* promoter region to confirm the integrity of the promoter region duplication and/or mutation. The single copy nature of *cyp51A* in the targeted integrants was confirmed using qRT-PCR, as described in (13).

### Drug Disc Diffusion Assay

Fresh spores of *A. fumigatus* were suspended in 1X phosphate-buffered saline (PBS) supplemented with 0.01% Tween 20 (1X PBST). The spore suspension was counted using a hemocytometer to determine the spore concentration. Spores were then appropriately diluted in 1X PBST. For the drug diffusion assay, ∼10^6^ spores were mixed with 10 ml soft agar (0.7%) and poured over 15 ml regular agar (1.5%) containing minimal medium. A sterile paper disk was placed on the center of the plate, and 10 μl of 2 mg/liter voriconazole was spotted onto the filter paper for analysis of wild-type and TR34 promoter mutant-containing strains. The same protocol was followed by 10 μl of 3.2 mg/liter voriconazole was spotted onto the filter disk for the analysis of the TR46 promoter-containing strains. The plates were incubated at 37°C and scanned after 72 hours.

### Western Blotting

Western blotting was performed as described in (13). The Cyp51A peptide polyclonal antibody used here has been detailed in the reference above, and was used at a 1:500 dilution.

### Measurement of mRNA level

Reverse transcription quantitative PCR (RT-qPCR) was performed as described in reference (13), with the following modification. The Ct value of the gene coding for *tef1* was used for normalization of variable cDNA levels to determine the fold difference in transcript levels with respect to *cyp51A*.

